# Use of a Sire MGS model to disentangle paternal and maternal origins of genetic variance in lifetime productivity of tropical dairy cattle

**DOI:** 10.64898/2026.03.09.710651

**Authors:** A. Menéndez-Buxadera

## Abstract

Data from 80,713 first-calving cows (1984 to1989) of the Holstein, Mambi, and Siboney breeds, belonging to seven large dairy enterprises in Cuba and progenies of 1,297 sires, were analyzed. For each cow, the average across all lactations for at least 14 years after first calving was defined as individual productivity (PI), and the corresponding lifetime sum as accumulated productivity (PA); both traits were. Two genetic models were fitted: a classical Animal Model (M1) and a Sire maternal grandsire model (Sire MGS; M2), aimed at partitioning additive genetic variance into paternal and maternal-line components. Heritability estimates under model M1 were moderate (h2 ≈ 0.135 to 0.140), whereas M2 yielded higher values (h2 ≈ 0.158 to 0.170), reflecting increased additive variance due to a better connectedness across herds. Using estimated breeding values (EBV) for PI and PA, a global cow merit index (H1) was defined under M1. Under M2, a parental index (IM2) combining four standardized predictors (paternal and maternal-grandsire EBV for PI and PA) was constructed. Multiple regression of H1 on IM2 showed that the paternal and maternal-grandsire paths accounted for 73% and 27% of the variation, respectively, indicating a non-negligible maternal-line contribution. Model M2 provided the best overall fit according to information criteria and cross validation using two independent subsamples and the full population yielded correlations of 0.870 to 0.881, demonstrating strong predictive ability and stability of IM2 rankings. These results support the Sire MGS model as a structural extension of the Animal Model for breeding programs targeting lifetime productivity in tropical dairy cattle.

## Introduction

The Animal Model (AM) has been the standard approach in modern quantitative genetics (Henderson, 1984), largely because it exploits all pedigree information to predict breeding values. However, AM typically aggregates sources of additive genetic variation into a single component (Meyer, 1992). This facilitates global estimation, but it does not distinguish between potentially different contributions from the paternal and maternal pathways. In most contemporary dairy systems, calves are artificially reared; thus, the maternal environmental effect can be assumed to be null or marginal, by contrast, maternal genetic transmission may still be relevant, yet evidence on its magnitude is limited. When both ancestral pathways contribute unequally and the statistical model does not reflect this structure, biologically informative variation can be lost (Willham, 1972) and variance–covariance components may be biased, thereby affecting the efficiency of breeding programs. Under such conditions, a Sire–maternal grandsire (Sire– MGS) model, viewed as an extension of AM, may be useful. Although this approach has received limited attention, it can quantify total additive genetic variance while also identifying the relative importance of the paternal and maternal-line components. It may reduce confounding among effects, requires lower computational burden, and according to Jenko et al. (2013) can provide a practical advantage over AM for routine dairy genetic evaluation.

Lifetime productivity has high economic relevance because it is closely linked to herd profitability and sustainability (Ducrocq, 1999). However, genetic improvement is challenging due to low heritability and the influence of multiple genetic and environmental factors (Forabosco et al., 2009). In practice, increased milk yield is embedded in selection goals, but questions remain on best way to quantify the results : milk yield per lactation versus lifetime production. In recent years, longevity studies have intensified (Hu et al., 2022), this is a threshold trait expressed over time and, despite the availability of powerful software (Ducrocq & Sölkner, 1998), there remains no full consensus on trait definition and reporting (Forabosco et al., 2009; Hu et al., 2021). One operational alternative is to evaluate lifetime milk production, which is a function of resilience, productive life, and sustained performance. Such measures align closely with farmer interest in retaining animals with higher lactation levels over longer periods.

This perspective may be particularly relevant in the tropics, where individual recording systems can be inconsistent across regions and pedigree information may be incomplete or unverified (Mrode et al., 2020). Here, a large historical dataset on lifetime milk productivity under tropical conditions—where the sire and maternal grandsire of each cow were known—was subjected to a powerful statistical analysis to estimate variance components and breeding values, with the aim of quantifying genetic variation that may be masked under the single aggregated variance in the AM. The objective of this study was to estimate (co)variance components using both a classical Animal Model and a Sire–MGS model, to maximize the utility of available information and to identify the origin of genetic variation.

## Materials and Methods

### Population and data

A historical database comprising 526,028 lactations recorded between 1984 and 2003 across the country was available. From this database, 80,713 cows of the Holstein, MambÍ, and Siboney breeds were selected, belonging to seven large dairy enterprises in Cuba daughters of 1,297 sires. Editing criteria required cows to be first calving with 24– 40 months of age and to have subsequent performance recorded for at least 14 years. The pedigree included the animal, its sire, and its maternal grandsire (MGS); the MGS was used as a proxy for the dam for all cows. The traits analyzed were: (i) Individual Productivity (PI), defined as the average 305-day milk yield across all lactations; and (ii) Accumulated Productivity (PA), defined as the sum of 305-day milk yield across the entire productive lifetime. For redaction convenience, sometime m305 is used as a synonym for PI and s305 for PA.

### Statistical models and variance partitioning

The data were analyzed using bivariate linear mixed models (for PI and PA) with the software Echidna (Gilmour, 2022). The models used were as follows:

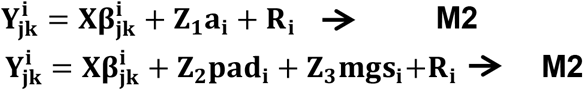

in which

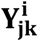 denotes the **jkth** observation of the **a**^**th**^ animal for the **i**^**th**^ trait (**i** = **X**_**1**_ = **PI; X**_**2**_ = **PA);** 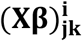 represents the fixed effects of contemporary groups defined by herd–year–season (20,758 levels), as well as the additive and heterosis covariates (**ad** and **he**) respectively). These coefficients were estimated according to the proposal of Akbas et al. (1993).

**a**_**i**_, **pad**_**i**_ and **mgs**_**i**_represent the random effects of the animal, sire and maternal grandsire, respectively.

**R**_**i**_ is a random residual effect common to all observations in each model.

**Z**_**1**_, **Z**_**2**_ and **Z**_**3**_ are incidence matrices linking the fixed and random effects to the data vector.

In these models it is assumed that:

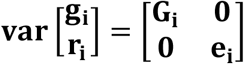

In which

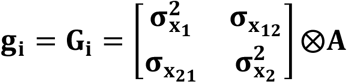 **⨂A** represents the matrix of variances 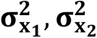 and genetic covariances **σ**_**x12**_between traits; ⊗ denotes the Kronecker product, and **A** is the numerator relationship matrix. The solutions for **g**_**i**_ with variance **G**_**i**_ may differ depending on which ancestor is included in **A (G**_**a**_, **G**_**pad**_ and **G**_**mgs**_ for animal, sire and mgs, respectively), allowing the origin and importance of the genetic (co)variance components attributable to the animal, its sire and its maternal grandsire to be distinguished.

On the other hand, the residual (co)variances are estimated as:

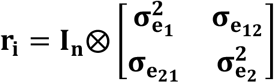

where **I**_**n**_ is an identity matrix of order equal to the number of observations, and

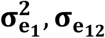, σ **e**_**12**_ are the corresponding residual (co)variances.

In this study, the same dataset and pedigree were used for all analyses, but the random effect was defined according to the ancestors considered. In **M1**, the traditional animal model approach was applied, estimating a combined variance as a single aggregated additive genetic component. Model **M2** allows the genetic variance to be decomposed by origin through the effects of the sire (pad) and the maternal grandsire (mgs), used as a proxy for maternal genetic effects. Echidna provides standard errors of the genetic parameters, as well as an output file with the breeding values (BV) and their standard errors, from which the prediction error variance (PEV) and the accuracy of the BV for each animal and for both traits were calculated. It should be emphasized that the variance biological interpretation may differ according to the structure of the statistical model used, whose particularities and formulations are presented in Table 1.

**Table 1.** General summary of genetic parameter estimation under both models^*^.

| Models | Parameters Estimated | Breeding Value and accuracy |
| --- | --- | --- |
| **M1: Aggregate variances**<br>$y_{jk} = X\beta_{jk} + Za + e$<br>$a \sim N(0, A \otimes G_a)$<br>$e \sim N(0, I \otimes e)$ | $\sigma_{ft}^2 = \sigma_a^2 + \sigma_e^2$<br>$h_a^2 = \frac{\sigma_a^2}{\sigma_{ft}^2}$ | $VG_a = \hat{a}$<br>$ACC_a = \sqrt{1 - \frac{PEV_a}{\sigma_a^2}}$ |
| **M2: Origin of variances**<br>$y_{jk} = X\beta_{jk} + Zs + Zm + e$<br>$s \sim N(0, A_{pad} \otimes G_{pad})$<br>$m \sim N(0, A_{mgs} \otimes G_{mgs})$<br>$e \sim N(0, I \otimes e)$ | $\sigma_{pad}^2 = 4\widehat{\sigma_{sire}^2}$<br>$\sigma_{mgs}^2 = 16\widehat{\sigma_{mgs}^2}$<br>$\sigma_{ft}^2 = \sigma_{pad}^2 + \sigma_{mgs}^2 + \widehat{\sigma_e^2}$<br>$h_{pad}^2 = \frac{\sigma_{pad}^2}{\sigma_{ft}^2}$<br>$h_{mgs}^2 = \frac{\sigma_{mgs}^2}{\sigma_{ft}^2}$<br>$h_{tot}^2 = \frac{\sigma_{pad}^2 + 0.5\sigma_{mgs}^2}{\sigma_{ft}^2}$ | $VG_{pad} = 2\hat{s}; VG_{mgs} = 4\hat{m};$<br>$VG_{tot} = VG_{pad} + VG_{mgs}$<br>$PEV_{tot} = 4PEV_{pad} + 16PEV_{mgs}$<br>$ACC_T = \sqrt{1 - \frac{PEV_{tot}}{\sigma_{pad}^2 + \sigma_{mgs}^2}}$ |

\*The meaning of the symbols in the text.

The results of both models were compared using classical information criteria: log-likelihood (LogL), Akaike information criterion (AIC) and Bayesian information criteria (BIC), to select the best-fitting model. To evaluate BV stability, parallel analyses were performed in two independent subsamples of cows defined according to alternating classes of the number of calving produced over time. Subsample 1 included cows with 1, 3, or 5 calvings (57% of data), whereas subsample 2 included cows with 2, 4, 6, 7, or 8 calvings (43%). BV from subsamples were compared with those from the full population using correlations.

### Breeding values and derived indices

For each model, individual additive breeding values (BV) were obtained for the traits individual lifetime productivity (m305) and accumulated lifetime productivity (s305). Under M1 (classical Animal Model), the BV were estimated at the animal level, 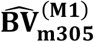 and,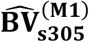 and were used as a reference for the available genetic variability. The overall genetic merit of animals under M1 was defined by means of an aggregated index H_1_, constructed from the breeding values of m305 and s305 following classical selection index theory (Hazel, 1943). For this purpose, standardized breeding values were computed for each trait as

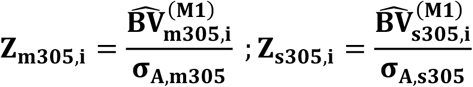

and the global index was then defined as

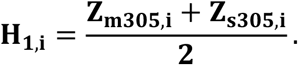

This index is equivalent to the aggregated genotype for lifetime milk productivity under Model M1 and was used as a benchmark to evaluate the predictive ability of the alternative modelling approaches.

Under Model M2, breeding values were obtained for each animal through both the paternal pathway and the maternal-grandsire pathway for the two traits:

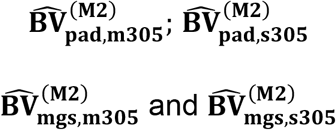

These four predictors were combined into a linear index I_M2_, representing the joint parental merit. Breeding values from M2 were expressed on a standardized scale, yielding for each animal the vector

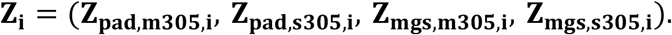

From this, an *i* × 4 matrix was constructed, which was used to compute the correlation matrix **R**_**pred**_ among these predictors and their correlations with the column vector H_1_. In this way, the correlation of each component of Z_*i*_with the daughters’ global index was obtained (**r**_**pred**,_**H**_ 1_).

Using these indicators and solving

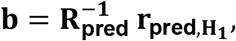

which is equivalent to fitting a multiple regression of H_1_on the four standardised breeding values for sire and maternal grandsire (Mrode, 2013), the resulting regression coefficients

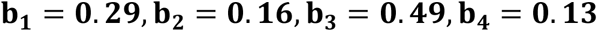

were used as weighting factors for the combined index **I**_**M2**,**i**_, defined as

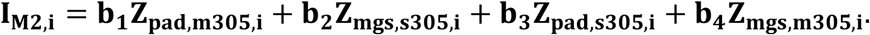

Thus, I_M2_ summarizes the weighted information from the sire and maternal-grandsire pathways as a joint contribution to the prediction of the global merit H_1_. Through this procedure, it is possible to differentiate the relative importance of the transmission pathway (sire vs. maternal grandsire) for each e trait (m305 vs. s305) in determining lifetime productivity.

## Results

All effects included in both models were significant (p < 0.0001) for both traits. Mean performance was 2336 ± 1152 kg for PI and 5733 ± 4896 kg for PA; cows produced 2.30 ± 1.4 lactations during the analyzed period. The **he** effects were significant for PI and the opposite for PA, while the **ad** effects were highly significant for both effects, representing a superiority of approximately 20-25% with respect to heterosis in PI and PA of both models. It is considered that no further details are necessary, these indicators have been shown only to illustrate the performance of the data set since the basic purpose of this study is the estimation of components of (co)variance.

Model comparison showed that M2 (Sire–MGS) had a higher likelihood than M1 (difference in -2logL = 19.6 units) and AIC decreased by 13.6 points in favour of M2, whereas BIC slightly favored M1. On the other hand, the validation results showed that the correlations between the indices obtained in each subsample and those estimated in the complete population were high: **r = 0.882** for sample 1 and **r = 0.870** for sample 2, which indicates a high stability of the genetic ranking in the face of substantial changes in the composition of the sample. According to these trends, it is considered that the choice of **M2** as the main model due to the increase in likelihood and the improvement of the AIC, together with the greater biological coherence, make it the most consistent option with the objectives of this work, about differentiating genetic variance in its paternal and maternal components.

Although both models were fitted with the same effects, the same data and the same pedigree, the results differed. For PI, the additive genetic variance was 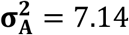 = 7.14 while the estimation of heritability 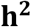 = **0. 141** ± **0. 05** in **M1**, whereas in M2 the total genetic variance was 10.6 and 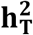 = **0. 170** ± **0. 055**. For PA, the patterns were similar, with 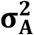= 7.811 and **h**^**2**^ = **0. 135** ± **0. 05** in M1, compared with **h**^**2**^ = **0. 158** ± **0. 05** in M2. For both traits, **M2** showed a better numerical fit, and the total heritability was higher than those from M1. All variance components and genetic parameters from both models are presented in Table 2.

**Table 2.** Variance components for the longevity measures PI and PA under both models.

| Individual Productivity |  |  |  |  |  |  |  |
| --- | --- | --- | --- | --- | --- | --- | --- |
| Models | $\sigma_a^2$ | $\sigma_m^2$ | $\sigma_a^2 + 0.5\sigma_m^2$ | $\sigma_T^2$ | $h_d^2$ | $h_m^2$ | $h_T^2$ |
| M1-Cow Total | 7.144 |  |  | 50.666 | 0.141 |  | 0.141 |
| M2- pad-mgs | 7.003 | 7.354 | 10.680 | 62.783 | 0.112 | 0.117 | **0.170** |
| Accumulated Productivity |  |  |  |  |  |  |  |
| Modelo | $\sigma_a^2$ | $\sigma_m^2$ | $\sigma_a^2 + 0.5\sigma_m^2$ | $\sigma_T^2$ | $h_d^2$ | $h_m^2$ | $h_T^2$ |
| M1-Cow Total | 7.811 |  |  | 57.692 | 0.135 |  | 0.135 |
| M2- pad-mgs | 7.727 | 6.593 | 11.024 | 69.551 | 0.111 | 0.095 | **0.158** |

Standard errors of heritability ranged from 0.006 to 0.008

The genetic correlations between m305 and s305 were very similar across models. In **M1, r**_**g**_ = 0.570 ± 0.03, whereas in **M2** were **r**_**g**_ = 0.578 ± 0.03and **r**_**g**_ = 0.501 ± 0.03 for the pad and mgs components respectively, indicating a positive genetic association of comparable magnitude between individual and accumulated productivity under both approaches.

The index **I**_**M2**_ = **0. 29**(**pad**_**s305**_) + **0. 16**(**mgs**_**s305**_) + **0. 49**(**pad**_**m305**_) + **0. 13**(**mgs**_**m305**_) summarizes the total genetic merit of sires for both traits. To facilitate its interpretation, the coefficients were summed and expressed relative to their total:

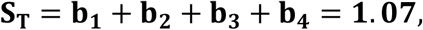

while the relative contributions of pad and mgs were

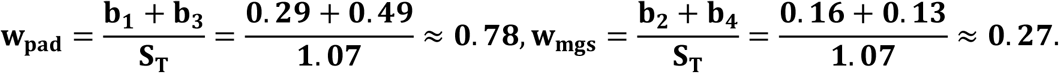

indicating that around 73% of the total influence comes through the paternal pathway, while the remaining 27% corresponds to the contribution of the maternal grandsire. In other words, approximately one quarter of the global genetic merit index of the daughters arises from the mgs pathway, which is theoretically expected, but the identification of this marginal contribution is by no means negligible.

The coherence between the parental merit summarized in **I**_**M2**_ and the merit expressed in the daughters was reflected in the regression of **H**_**1**_on this index: the estimated regression coefficients was **b** = **0. 481** ± **0. 001**, i.e. a value very close to the 0.5 expected under an additive model, in which approximately half of the parental merit is expressed in the offspring. This result indicates that the pad–mgs model consistently reconstructs the level of merit observed in the daughters, while simultaneously exploiting information from both the paternal and maternal pathways. This can be regarded as an important additional benefice of the pad–mgs model when compared with the classical Animal Model, which combines both pathways into a single aggregated additive effect.

The variability observed in both traits reveals a wide range of possibilities; in particular, the sire pathway may have the greatest impact given the inherent capacity of bulls to produce many progenies (Figure 1).

**Figure 1.**
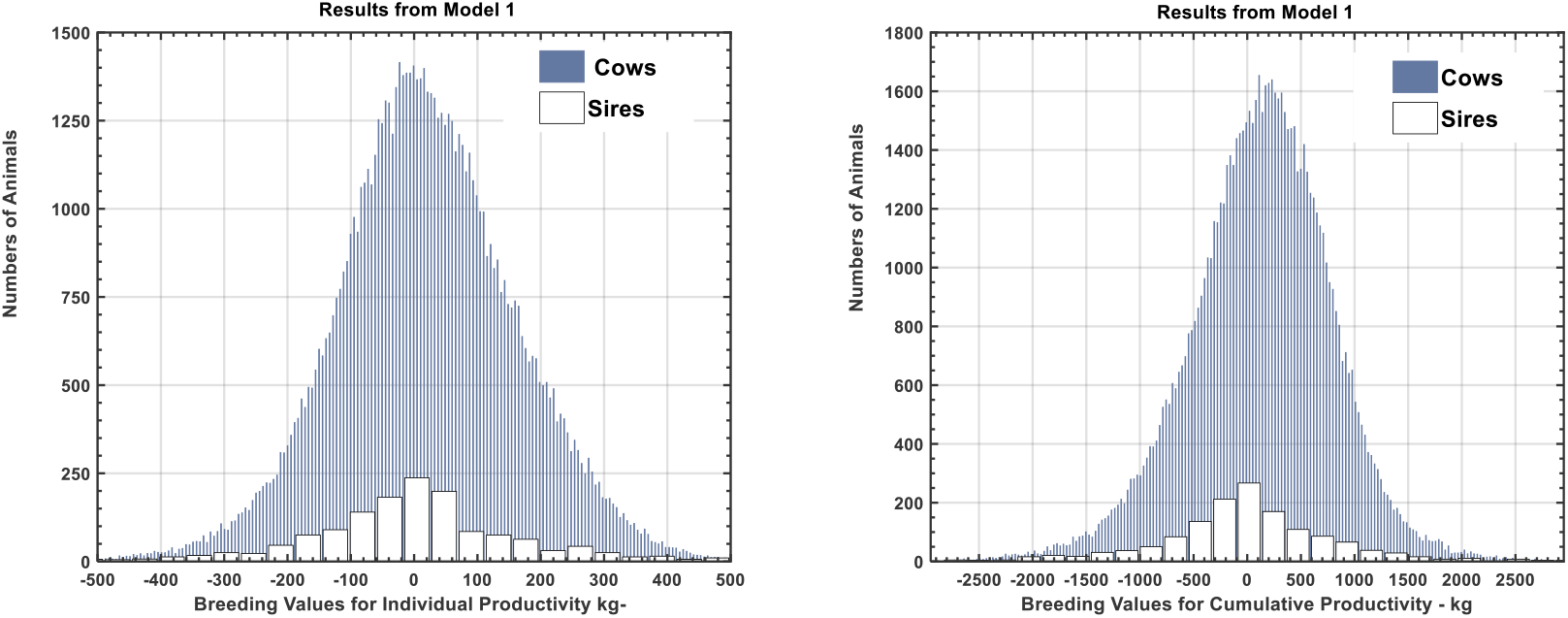
Frequency distribution of the Genetic Values of Individual and Cumulative Productivity according to the classical model.

However, the BV for this dataset has been estimated using different models and transmission pathways, generating a large volume of information that requires an integrated assessment, for which principal component analysis (PCA) is recommended. The PCA is a statistical procedure that makes it possible to summarize most of the information from multiple genetic predictors by minimizing the deviation of each one from an underlying line (eigenvector), without appreciable loss of the original variances. Figure 2 shows the results of this PCA applied to the BV for both traits as estimated by each model, together with the index **I**_**M2**_ presented in the previous paragraphs. The results indicate that between 96% and 98% of the variation is explained by the first eigenvector, leaving very little importance to the second and third eigenvectors. Note the distribution and vector directions in the PCA applied to the genetic values of PI and PA obtained with the classical model (M1) and the pad–mgs model (M2), together with the global index **I**_**M2**_ they follow the same pattern and are therefore robust estimators. Moreover, a single vector corresponding to the global genetic index **I**_**M2**_ by itself represents and explains almost all the existing genetic variance.

**Figure 2.**
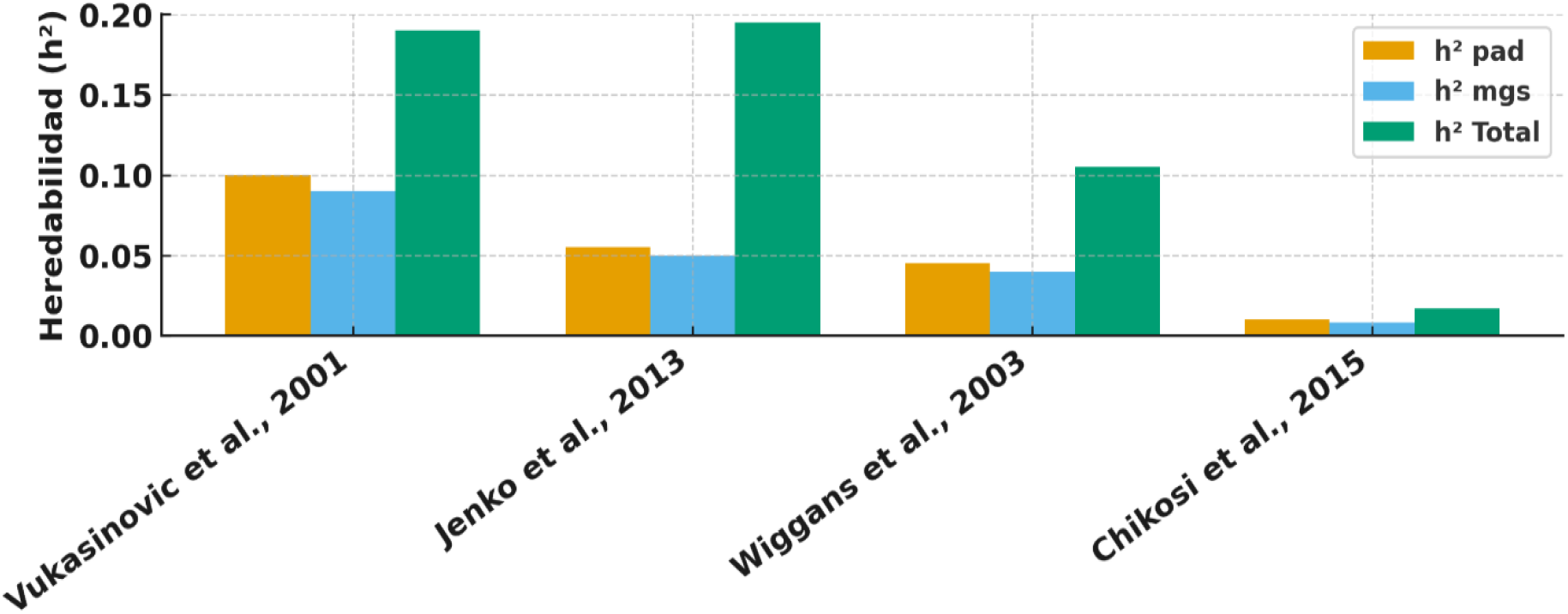
Representation of heritability estimates obtained through different pathways using the Sire–MGS model.

Note that the pad–mgs model (M2) not only increases the total genetic variance and heritability already shown in Table 2, but also organizes the genetic variation more efficiently, concentrating it along a main axis that can be exploited for evaluation.

Figure2. Vector distribution of the principal component analysis (PCA)^*^ applied to the breeding values for PI and PA obtained with the classical model (**M1**) and the pad–mgs model (**M2**), together with the global index

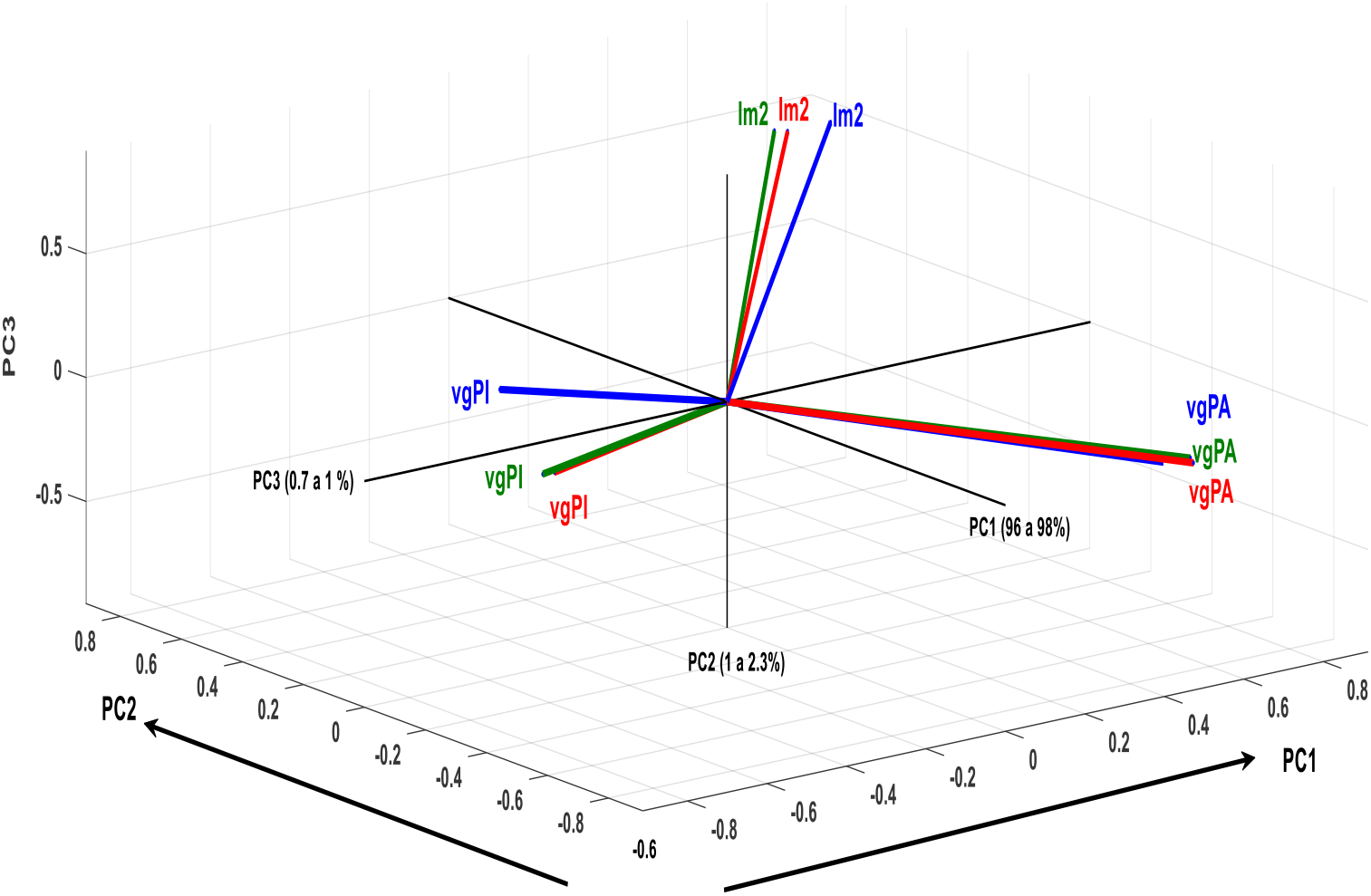
^*^Im2 represents the overall index of all sires for both variables (in green); in red it refers only to the parents of cows according to model 1, in blue it refers to parents of cows from model 2. vgPi and vgPA mean the same thing but PI and PA

## Discussion

In this article, the criteria of Individual Productivity and Accumulated Productivity were used to express lifetime milk output in a simple manner closely aligned with the practical reality of dairy production in Latin America, where production systems require animals that deliver a high total volume of milk, measured in kilograms. This approach is not intended to replace what is commonly referred to as longevity in animal science, a concept for which multiple definitions exist. In an excellent review, Hu et al. (2021) examined ten definitions of longevity used in sixteen countries and showed that, although there is consistency in reporting low heritability estimates, there is little uniformity in how this trait is defined and recorded. Taken together, this lack of standardization represents a serious obstacle to implementing more successful breeding programs.

Although the traits analyzed and the structure of the data may differ across studies, it is evident that the population represented here exhibited lower productivity than that reported for Costa Rica (Vargas-Leyton et al., 2024) and Honduras (Copas-Medina et al., 2023). These results were expected, according to the extensive analysis published by Hernández et al. (2018) using a very similar dataset under Cuban conditions. Regardless of this reality, greater attention to the importance of lifetime productivity is imperative, and the results of the present study demonstrate that there is ample scope to identify animals with high total lifetime production, whose economic relevance is unquestionable.

Heritability estimates for PI and PA obtained with the classical Animal Model (M1) were in the range of **h2 ≈ 0.135 – 0.140** (Table 2), values consistent with those Hernández et al., (2011) in Cuba reported (h2 ≈ 0.05–0.15;), Honduras (h2 ≈ 0.15–0.25); Copas-Medina et al., 2023), and Mexico (h2 ≈ 0.13–0.15; AbadÍa-Rojas et al., 2016). At the same time, the estimates from this study lie at the upper limit of the 24 values summarized by Hu et al. (2021). To our knowledge, systematic and periodic studies on this topic are scarce in Latin America, probably due to limitations in recording systems and mostly to an underestimation of the economic importance of these traits, an issue that clearly deserves greater attention.

The differences observed in total heritability in Table 2 can be explained using **sire (pad)** and **maternal grandsire (mgs)** as random effects, which substantially improves genetic connectedness across the data compared with M1. Through this pathway, ancestors tend to have offspring more widely distributed across numerous herds and over time. This feature of Model M2 provides greater efficiency in the estimation of the genetic component by reducing potential confounding with contemporary group effects and other environmental factors, thereby increasing the accuracy of breeding values. This better connectedness, as discussed by Quaas and Pollak (1980) and Mrode (2014), results in a more compact diagonal and better off-diagonal representation of the elements of the inverse relationship matrix A^-1^, providing the data structure required for algorithms such as REML to yield more precise estimates of genetic parameters and improved predictive ability for daughters.

In terms of breeding value prediction, the validation analysis produced correlations ranging from **0.870 to 0.881** between subsamples 1 and 2 and the global genetic index I_M2_. These values fall within the range commonly reported in the literature for studies assessing the stability of breeding values in complex, low-heritability traits such as longevity and fertility, where correlations of **0.80–0.90** are considered indicative as high degree of ranking consistency (Ducrocq et al., 1988; Boichard et al., 1995). These advantages are particularly relevant in populations with limited recording systems, where Model M2 offers a practical means of maintaining precision and connectedness at an acceptable computational cost, thereby justifying a re-evaluation of current breeding strategies.

The use of a **Sire–MGS model** is often criticized because of the possible covariance between direct and maternal effects; in the present study, this covariance was assumed to be zero. Maternal influence has two components: one is an environmental effect, related to the care provided by the dam, whose importance is largely eliminated under artificial calf-rearing systems, and the second is a **maternal genetic component**, which may play an important role but has received limited attention. In Model **M1**, an aggregated variance derived from both ancestors is estimated, whereas in **M2** it is possible to differentiate its origin. As shown in Table 2, the additional component identified is by no means negligible.

Figure 2 summarizes results from several studies applying Sire–MGS models and shows a consistent pattern in which this statistical approach leads to increases in total heritability.

It can be noted that the MGS and sire pathways contribute very similarly to each reference, and that when they are combined, total heritability increases substantially, leading to greater reliability of the estimated breeding values. The same tendency was observed in the present study. Consequently, the Sire–MGS model may be regarded as a valuable auxiliary tool to the Animal Model, capable of providing more precise and less biased estimates in scenarios where individual recording systems are poorly developed and environmental conditions are highly heterogeneous, as is often the case in tropical production systems.

To conclude this section on the potential benefits of the Sire–MGS approach, a simple simulation based on the results in Table 2 was carried out to compare the expected genetic gain from selecting the top 5% of sires. The results are shown in Figure 3. The overall trend indicates that, under any of the scenarios considered, the Sire–MGS model can provide an advantage of approximately **10–17%** over the classical Model 1. Achieving better results with the same amount of information is equivalent to a more efficient management of the breeding program.

**Figure 3.**
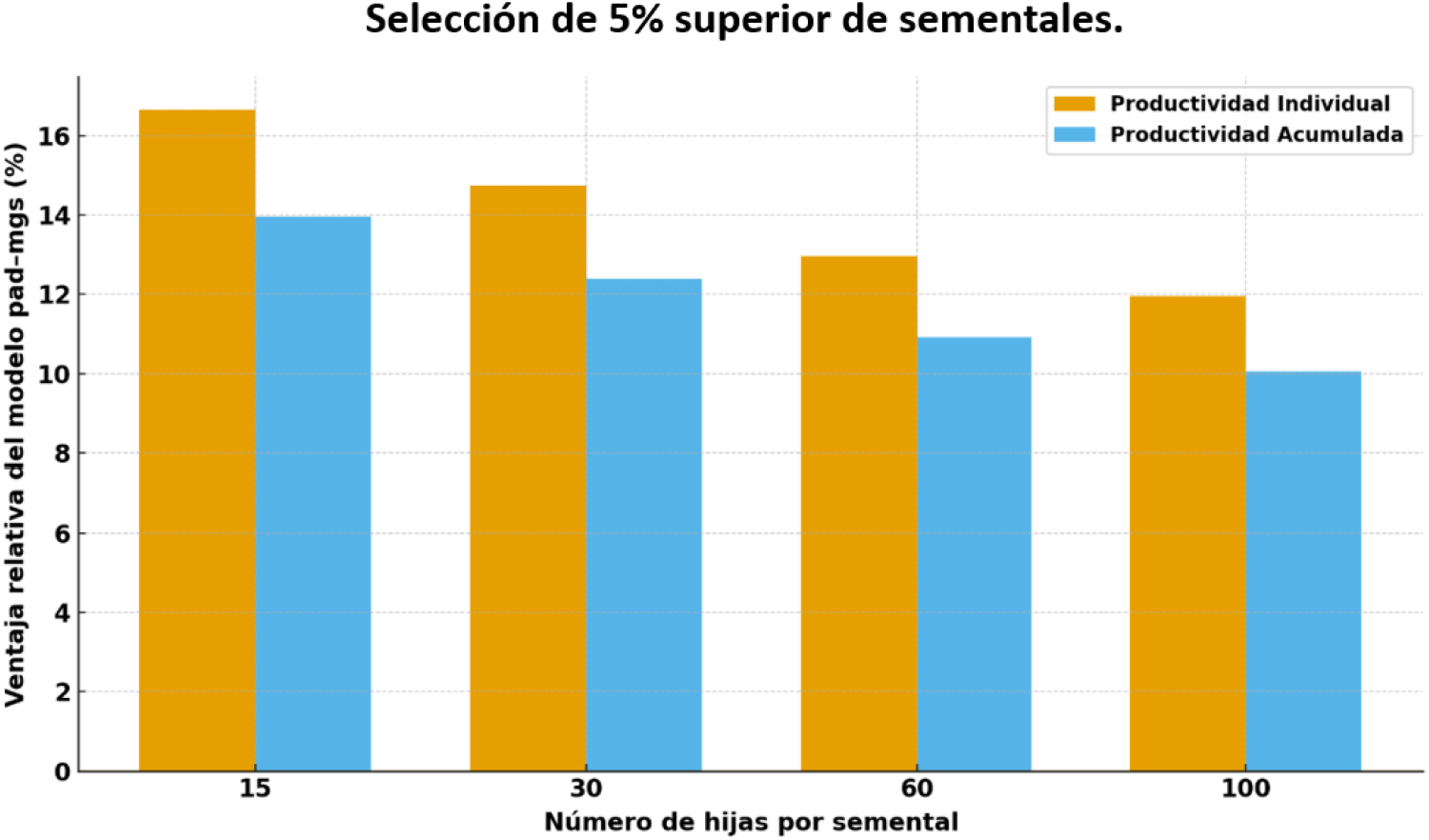
Progreso genético relativo del pad-mgs frente al modelo animal(M1)seleccióde 5%superior de sementales.

## Conclusion

The Sire–MGS model can be regard as a complement of the classical Animal Model by allowing additive genetic variance in lifetime productivity to be partitioned into paternal and maternal-line origins. Under the present data, this partition was associated with a 17– 20% increase in total heritability for PI and PA relative to the aggregated Animal Model and supported an index (**I**_**M2**_) in which approximately 73% of the parental contribution was attributed to the sire pathway and about 27% to the maternal-grandsire pathway. These findings support reconsideration of evaluation strategies for tropical dairy populations, particularly where recording and connectedness are limiting factors.

This study shows that the pad–mgs model is a very promising alternative to the classical Animal Model, as it allows the additive genetic variance to be decomposed into its paternal and maternal origins. Under the conditions of the database analyzed here, this decomposition translated into an increase of 17–20% in the total heritability of the longevity/productivity complex relative to the animal model, and also made it possible to align, organize and concentrate the genetic variation more efficiently along a main axis that is easily interpretable and directly exploitable for evaluation. Taken together, these features open the door to achieving greater responses to selection for these traits.

The construction of the parental index **I**_**M2**_ made it possible to quantify the relative contribution of each pathway, showing that approximately 73% of the aggregate genetic merit of pad and mgs is channelled through the sire pathway and around 27% through the maternal grandsire. Far from being marginal, this maternal fraction represents nearly one quarter of the usable genetic merit and demonstrates that the maternal effect (captured here via the mgs proxy) has been underestimated when working solely with a single aggregated variance in the Animal Model.

Additionally, the use of pad and mgs as random effects improves connectedness across herds and time periods, reduces the risk of biases associated with confounding with contemporary group effects or potential genotype–environment interactions, and strengthens the structure of the matrix **A**^-**1**^, favouring more stable and faster convergence of REML algorithms. Although this study focused on longevity, the results suggest that the pad–mgs model could be equally useful for other economically important traits in which the maternal pathway plays a relevant role; our preliminary analysis of first-lactation milk yield points in the same direction. Overall, these findings support a reconsideration of genetic evaluation strategies, especially in tropical populations, by explicitly incorporating the contribution of paternal and maternal lines into breeding programs.

Finally, it should be borne in mind that the mgs previously occupied the position of sire, but in the dynamics of a selection program only sires with the highest BV go on to become maternal grandsires. Thus, this pathway represents a selected subset of bulls with superior genetic merit. Discussing this feature in depth lies beyond the scope of the present article, but it clearly deserves attention in future work.

## Acknowledgements

I express my gratitude to MSc José Raúl Pérez González, Universidad Politécnica Territorial de Maracaibo, Venezuela, for his ongoing discussions and support during the development of this study.

